# scTPA: A web tool for single-cell transcriptome analysis of pathway activation signatures

**DOI:** 10.1101/2020.01.15.907592

**Authors:** Yan Zhang, Yaru Zhang, Jun Hu, Ji Zhang, Fangjie Guo, Meng Zhou, Guijun Zhang, Fulong Yu, Jianzhong Su

## Abstract

The most fundamental challenge in current single-cell RNA-seq data analysis is functional interpretation and annotation of cell clusters. The biological pathways in distinct cell types have different activation patterns, which facilitates understanding cell functions in single-cell transcriptomics. However, no effective web tool has been implemented for single-cell transcriptomic data analysis based on prior biological pathway knowledge. Here, we introduce scTPA (http://sctpa.bio-data.cn/sctpa), which is a web-based platform providing pathway-based analysis of single-cell RNA-seq data in human and mouse. scTPA incorporates four widely-used gene set enrichment methods to estimate the pathway activation scores of single cells based on a collection of available biological pathways with different functional and taxonomic classifications. The clustering analysis and cell-type-specific activation pathway identification were provided for the functional interpretation of cell types from pathway-oriented perspective. An intuitive interface allows users to conveniently visualize and download single-cell pathway signatures. Together, scTPA is a comprehensive tool to identify pathway activation signatures for dissecting single cell heterogeneity.

## INTRODUCTION

Single-cell RNA sequencing (scRNA-seq) technology has been widely used to characterize cell-to-cell heterogeneity (1). The single-cell transcriptome analyses uncover the new and unexpected biological discoveries compared to traditional “bulk” cell methods (2). Many computational methods have been developed for cell clustering, marker genes identification and visualization of single-cell RNA-seq data (3, 4). However, functional interpretation of cell clustering remains challenge for scRNA-seq data analysis.

Pathways are biological network models defining how biomolecules cooperate to accomplish specific cellular functions in distinct cell types, which are crucial to disease subtype classification (5), functional annotation of cellular diversity (6) and drug discovery (7). In single-cell studies, pathway activation analysis has become a powerful approach for extracting biologically relevant signatures to uncover the potential mechanisms of cell heterogeneity and dysfunction in human diseases (8, 9). For example, the pathway signatures exhibit the significant activation difference in breast cancer (10) and Alzheimer’s disease cells (11) using gene set enrichment analysis. However, there is a lack of online web server for the comprehensive analysis and visualization of single-cell transcriptome data based on prior biological pathway knowledge.

Here, we developed scTPA (http://sctpa.bio-data.cn/sctpa), which is a web-based platform dedicated to pathway signature discovery and functional interpretation of scRNA-seq data in human and mouse. Abundance of high-quality curated biological pathways with different functional and taxonomic classifications were manually collected, facilitating the pathways selection according to the research context and interests. scTPA incorporates four widely-used methods to calculate the pathway activation profiles and provides the flexible parameters for downstream analysis. Based on well-known biological pathways or user-defined pathways, clustering analysis and cell-type-specific activation pathway identification were performed, which allow a better understanding of their potential functions from pathway-oriented perspective. The scTPA provides an easy-to-use interface for viewing and download of pathway activity scores, cell clustering, pathway signatures and the associated gene expression.

## WORKFLOW OF scTPA

The scTPA is a web tool for single-cell transcriptome analysis and annotation based on pathway activation signatures in human and mouse. Firstly, user could upload the single-cell gene expression profiles to scTPA for data normalization, filtration and imputation. According to user’s interests, biological pathways would be selected from our collected pathway library of different function and taxonomy or user defined pathways. Secondly, four widely-used enrichment analysis methods are provided to rapidly compute pathway activity score (PAS) of each cell. Cell types could be determined optionally by users or defined by clustering analysis based on PAS matrix. Finally, statistical analysis is performed to identify the cell-type-specific activation pathways (CTSAPs), which allows a better understanding of cell type and biological status. Multiple interactive visualizations of outputs are also provided. The detailed schematic view is showed in Figure 1.

**Figure 1.**
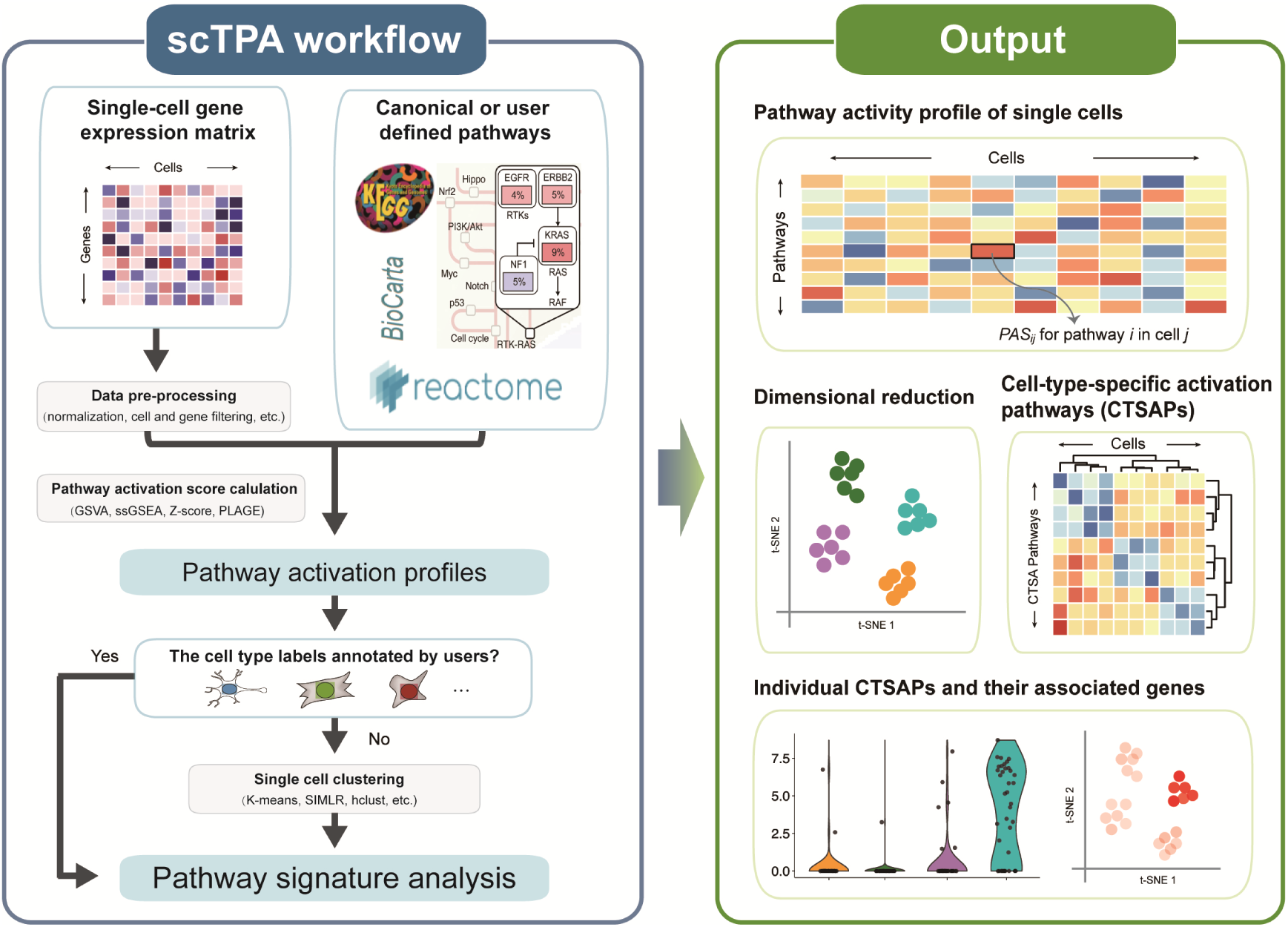
An overview of scTPA web tool. Input single-cell gene expression profile was pre-processed and the PAS matrix would be calculated based on interested pathway signatures using the enrichment-based methods. The cell-type label could be provided either by the user or unsupervised clustering analysis of PAS matrix, and the pathway signature analysis was performed. The resulting data for single cell transcriptome dimension reduction, clustering, pathway signatures identification, visualization and download.

## INPUT&DATA PROCESSION

### scRNA-seq profile

The required input data of scTPA is a processed single-cell gene expression matrix where columns correspond to cells and rows correspond to genes. The input file is read count or RPKM/FPKM/TPM/CPM of single cells generating from different platforms such as 10X genomics and Smart-seq, etc. For users’ convenience, scTPA supports the data upload of the input file with a pre-compression.

Multiple data normalization methods are provided including log transformation, quantile normalization and Z-score. Users can also choose to remove poor cells and genes that are not detected with enough proportion. The single cell profile contains excess zero or near zero counts due to extensively dropout events caused by the low amounts of mRNA sequenced within individual cells. The scTPA also provides option to impute the missing values(12).And scTPA only imputated genes with dropout (i.e. expression equal to zero) rates larger than 50% to avoid over-imputation.

### Biological pathways

To facilitate the evaluation of pathway activation at single-cell resolution, the ‘Canonical pathway’ and ‘Extended pathway’ options were provided for the user to select users interested literature-curated pathways.

Canonical pathways of scTPA contain 51,210 human and 1,762 mouse pathways from seven widely used pathway databases including BioCarta, HumanCyc (13), KEGG (14), PANTHER (15), PharmGKB (16), Reactome (17), SMPDB (18), which were retrieved from R package graphite (19). Notably, these literature-curated pathways were grouped into 6 different catalogs including general pathways, metabolic pathways, signaling and regulatory pathways, genome maintenance pathways, drug & small molecules pathways and cancer pathways (20). It facilitates selecting the relevant pathway database suitable for researchers interested context.

Extended pathways of scTPA represent many functional sets of unordered and unstructured collections of genes, which were associated with a specific biological process, genomic location, disease, cell identity, cell state or cell fate. They have more wide coverage of biological functions with genetic and chemical perturbation, computational analysis of genomic information, and additional biological annotation relatively to traditional canonical pathways. The currently extended pathways contain 19,367 pathways from 9 categories for human, and 19,385 pathways from 5 categories for mouse, respectively. They were collected from The Molecular Signatures Database (MSigDB V7.0) (21) and Gene Set Knowledgebase (GSKB) (22). In addition, the user could simultaneously upload their interested pathways, which are not cataloged by scTPA for specific scRNA-seq analysis.

### PAS calculation

Four classic methods including ssGSEA (single sample gene set enrichment analysis), GSVA (gene set variation analysis), PLAGE (pathway level analysis of gene expression) and Z-scores were incorporated into scTPA to measure the activation of pathways signatures for single cell transcriptomes, respectively. These methods generally calculate the enrichment scores with statistically significance from the expression-level rank statistics for a given pathway using the improved the R/Bioconductor package GSVA (23). To increase the computation efficiency, we rewrote the main loop function of GSVA which could achieve a 1.4-56 fold decrease for the runtime of massive parallel scoring pathway activation from processed gene expression matrix (supplementary information). This is a desirable feature for fast calculation of PAS, as the necessary in the analysis of single-cell gene expression data with large cell number.

### Unsupervised cluster analysis

Unsupervised cluster analysis is a useful exploratory tool to dissect the heterogeneity of complex populations. If the cell-type label file not pre-defined by the user, scTPA provided six different clustering methods (24, 25) including Seurat, K-means, K-mediods, SIMLR, DBSCAN and hclust to cluster cells based on PAS matrix. Main clustering parameters such as number of clusters, resolution, number of neighbors, dimensions of PCA were provided. Cell type annotation was further used for the following pathway signature analysis.

### Identification of cell-type-specific activation pathways

Pathways signatures are important for unveiling and characterizing the cell types and their functional states. Based on the PAS matrix of individual cells, five different statistical methods such as nonparametric Wilcox on rank sum test and likelihood-ratio test, and fold-change analysis were provided for CTSAPs identification. They may distinguish cell populations into the case-control groups consisted of the interested cell type and all other cells. scTPA can help the user find CTSAPs that are statistically significant activated among different cell types determined by the users or clustering analysis.

## OUTPUT

After the user submits the input data, a new tab is automatically opened to display the job progress of scTPA analysis. All the resulting files are available for download to users directly in the same page when the job is completed. Typically, the text-based files of pathway activation score (PAS) matrix, cell-type labels and statistics of pathway signatures and associated gene expression can be downloaded via “download” buttons or the corresponding web plugins. The web tool also provides the figures for visualization of PAS matrix, dimensional reduction and cluster analysis and pathway signatures. Specifically, a heatmap plot for the entire PAS matrix of single cells from a global view was provided. Interactive plots in 2D and 3D for dimensionality reduction generated using methods of t-distributed stochastic neighborhood embedding (t-SNE) (26) and Uniform Manifold Approximation and Projection (UMAP) (27) were provided to visualize the differences between cell populations. In addition, the statistical results and visualization of pathway signature analysis were provided. A heatmap plot was used to show the PAS profile of the significantly activated pathway signatures in each cell type. For each CTSAP in the corresponding cell type, we provide a UMAP plot and a box plot to display the PAS distributions across different cell types. We also provide the heatmap for gene expression in the CTSAPs to explore how the transcriptional changes affect the pathway activation of the various cell types.

## CASE STUDIES

To illustrate the function and utility of scTPA, we applied a processed data set of gene expression profiles for melanoma study (GEO accession number GSE72056, including 4054 cells) (28), which covered a variety of non-malignant cell types including B cells, T cells, macrophage, endothelial cells, cancer-associated fibroblasts (CAFs) and natural killer cells (NK), and malignant tumor cells.

### Case 1

To perform this analysis, we uploaded the single-cell gene expression TPM and the corresponding cell-type label files to the web server. Owing to the missing values in single-cell profiles were more than 80 percentage, we used the parameter for missing value imputation and started scTPA analyses with the other parameters follow the default setting.

The PAS matrix of single cells was calculated based on KEGG pathways using GSVA method, and its global view was showed in a heatmap (Figure 2A). The dimensionality reduction of PAS matrix with UMAP method showed in the 2D plot (Figure 2B). We found different cell types could be significantly distinguished based on pathway signatures, consistent with the original study (28).

**Figure 2.**
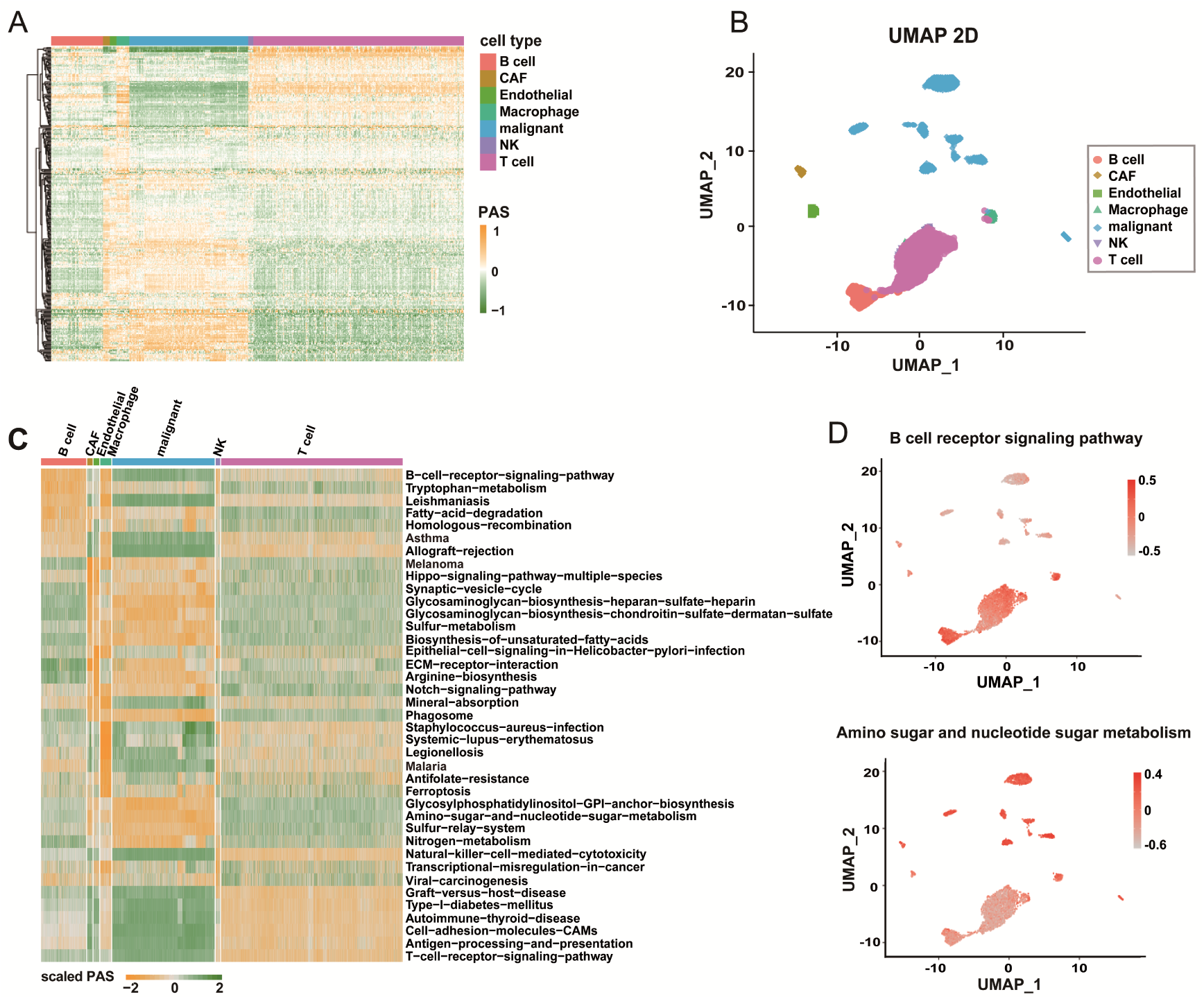
The analysis of melanoma dataset using scTPA. (A) The heatmap shows the pathway activity profile of highly variable pathways in the scRNA-seq data. Rows represent pathways, columns represent single cells. (B) UMAP-dimension reduction of cell populations based on the PAS profile. (C) The heatmap plot shows significant CTSAPs in each cell type. The colors represent user-defined cell type labels. (D) Visualization of individual activation pathway with UMAP plot.

Using the likelihood-ratio test, 185 out of 293 KEGG pathways were identified as CTSAPs using threshold of adjust P-value < 0.01 and average fold change > 0.25, and the 50 significant CTSAPs were displayed in the heatmap (Figure 2C). In different cell subpopulations, we found CTSAPs are closely related to the corresponding cell identities and their functional states. For example, B cell receptor signaling pathway and T cell receptor signaling pathway were markedly activated in B cells (P-value = 1.02e-214) and T cells (P-value = 1e-216), respectively. Allograft rejection pathway was simultaneously across immune cells including B cell (P-value = 6.13e-25), T cell (P-value = 9.47e-278) and macrophage cell (P-value = 3.27e-44), which was completely inactivated in tumor associated cell types of malignant and CAF cells. Natural killer cell mediated cytotoxicity pathway (P-value = 4.00e-26) and Platelet activation pathway (P-value = 8.67e-28) were specially activated in NK cell type, which are crucial for cellular immune defense mediated by NK cell.

Of note, we found that malignant cells exhibit a common pattern of global up-regulation of activities of metabolic pathways comparing to non-malignant cells. Nine of top 10 melanoma-specific activated pathways, such as Glycosaminogly can biosynthesis and Sulfur metabolism, were metabolic pathways reflecting different aspects of cellular metabolism. Our findings with scTPA analysis of single-cell gene expression profiles provide a global picture of pathway signatures for individual cells, which could provide new insight for annotation and understanding of cell types and their functional states on the basis of their preferentially or distinctively activated pathway signatures.

### Case 2

Next, we attempted to test whether scTPA could potentially dissect the heterogeneity of tumor cell population and reveal the potential cell subpopulations from the pathway-oriented view.

We extracted the gene expression profiles of malignant cells from the melanoma dataset and selected Hallmark gene sets of human cancer as pathway signatures to perform scTPA analysis. The malignant cells could be clearly classified into 8 groups based on 50 classic cancer hallmark pathways using unsupervised clustering method Seurat. Although cancer hallmarks are general features for different tumor cells, the dimensionality reduction analysis also demonstrated that the malignant cells of different types are clearly separated from one another (Figure 3A). The CTSAPs were identified for each cell subpopulations and the functional interpretation of the cell populations derived from PAS-based classification (Figure 3B). We found some hallmark pathways were simultaneously activated in multiple cell clusters, such as Angiogenesis in C2 and C8, Oxidative phosphorylation in C7 and C8, G2M checkpoint in C3, C4 and C7. In the cell cluster C1, we found several hallmark pathways, including inflammatory response, Hedgehog signaling and Interferon alpha response, were exclusively activated (Figure 3C). User could also inspect expression patterns of genes in CTSAPs for cell clusters of interest (Figure 3D). Our present analysis indicates that quantifying variation in oncogenic signaling pathways of individual malignant cells could explain the underlying mechanism driving tumor cell identity and functional states.

**Figure 3.**
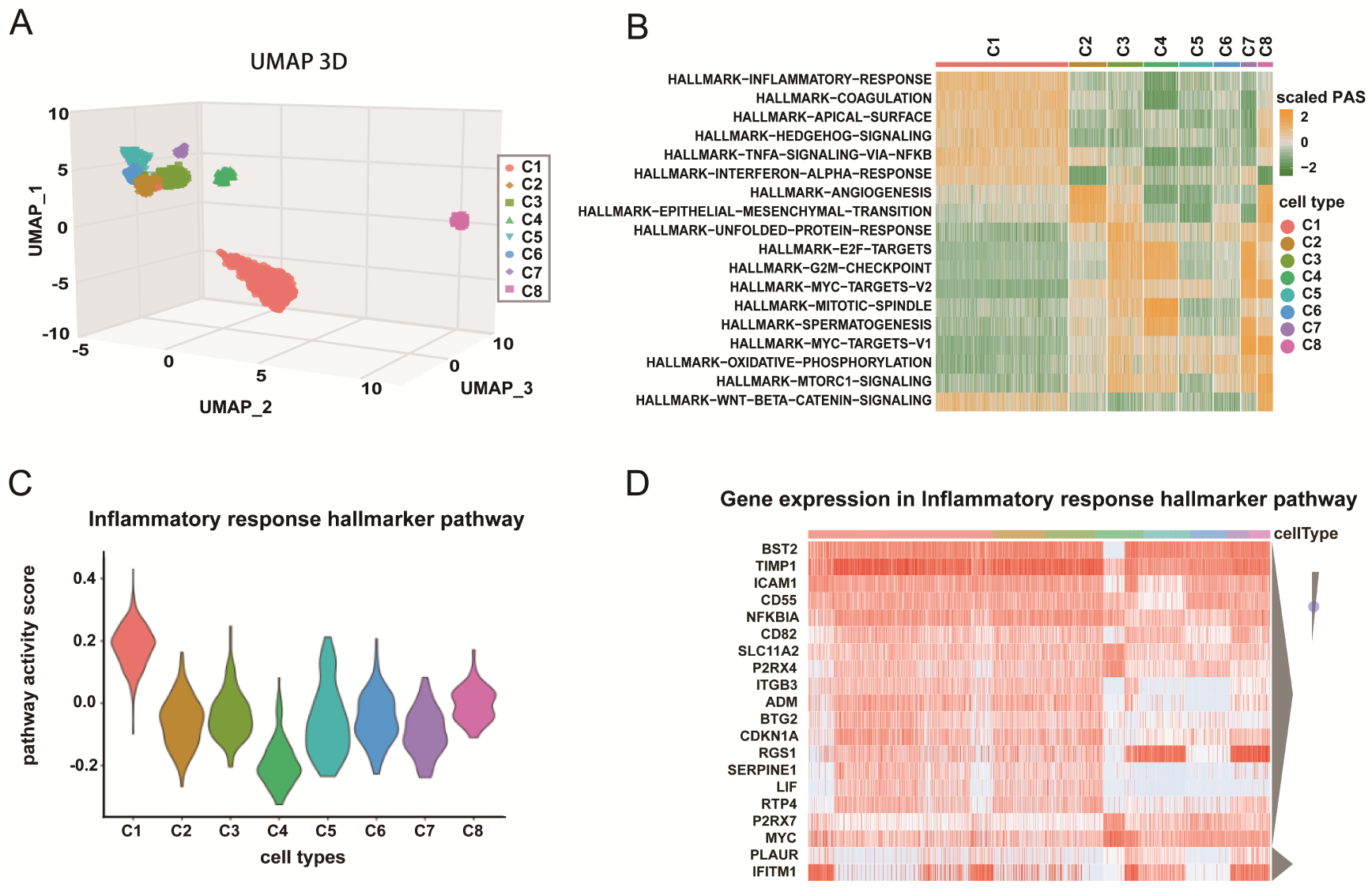
The analysis of single-cell transcriptome of malignant cells from melanoma dataset. (A) UMAP plot displaying all malignant cells, where each cell is clustered into one of the 8 clusters (distinguished by their colors). (B) The conserved cell-type-specific pathway signatures. The heatmap displays the PAS profile. Rows represent pathway signatures. Columns represent single cells, color-coded by their cell types. (C) Pathway activation distribution of a user-interested pathway signature. (D) The heatmap displays the gene expression patterns of the selected cell-type-specific activation pathway.

Overall, these two cases demonstrated that scTPA could not only classify, annotate and interpret the functional outcomes for single cell transcriptome data with known cell types, but also enable potentially deciphering the heterogeneity complexity of cell populations with unknown cell types.

## SUMMARY AND FUTURE DEVELOPMENTS

We developed an open-access, user-friendly web-based tool, scTPA, which provides a one-stop shop for single cell transcriptome dimension reduction, clustering and visualization by quickly evaluating the activation of biological relevant pathways. By identifying significantly activated pathways, scTPA is able to uncover biologically relevant subpopulations and further provides new insights for dissecting the complex heterogeneity of unlabeled cell subpopulations. The tool will be continuously updated and improved in the future to make it easily accessible through a web interface for in-depth single-cell transcriptome analyses. It is necessary to integrate more valid tools for clarifying the interplay between cell types and functional states in space and time. Topological information of the structure of connections among genes in the pathways should be taken into consideration to enhance estimation of pathway activation. In addition, combined machine learning approaches with literature-based knowledge could discover more meaningful pathway signatures in the future, which may be useful for annotation and interpretation of single-cell transcriptomes.

## DESIGN AND IMPLEMENTATION

The internal programs of scTPA are implemented using BASH, C++, PYTHON, JavaScript, MySQL, and R scripts. The online visualization was implemented using Highcharts (https://www.highcharts.com/), d3 (https://d3js.org/) and R package Seurat (24). Our system deployed on a server with 64 GB of RAM and sixteen 2.6 GHz Xeon CPUs.

## AVAILABILITY

http://sctpa.bio-data.cn/sctpa—this website is free and open to all users, it can be accessed by any major modern browsers such as Google Chrome, Mozilla Firefox and Safari.

